# Spontaneous formation of different forms of alpha-synuclein fibrils from a recombinant protein

**DOI:** 10.1101/2024.10.15.617481

**Authors:** V.V. Egorov, N.A. Grudinina, D.S. Polyakov, Y.A. Zabrodskaya, N.V. Gavrilova, M.M. Shavlovsky

**Affiliations:** Federal State Budgetary Scientific Institution ‘Institute of Experimental Medicine’, 197022, Akademika Pavlova street 12, St. Petersburg, Russia; Smorodintsev Research Institute of Influenza, Russian Ministry of Health, 197376, Prof. Popov St. 15/17, St. Petersburg, Russia; Institute of Biomedical Systems and Biotechnology, Peter the Great St. Petersburg Polytechnic University, 194064, Politekhnicheskaya 29, St. Petersburg, Russia; Federal State Budgetary Educational Institution of Higher Professional Education “Saint-Petersburg State University”, 199034, Universitetskaya embankment, 7-9, St. Petersburg, Russia

## Abstract

Alpha-synuclein is a protein, the conformational changes of which lead to the development of such socially significant diseases as Parkinson’s disease and amyotrophic lateral sclerosis. The methods for differential diagnostics of these diseases based on the use of alpha-synuclein in a non-native conformation obtained from patients as a seed for inducing fibrillogenesis and studying the morphology of the resulting amyloid-like fibrils were described in a number of studies. The authors associate such properties of the seed with the presence of post-translational modifications in the protein obtained from patients. At the same time, the production of fibrils differing in morphology from recombinant alpha-synuclein under various conditions of fibrillogenesis is also described. In this work, we show that the formation of morphologically distinct fibril types from recombinant alpha-synuclein lacking post-translational modifications is possible under the same conditions, and that spontaneously arising different fibril types, when used as a seed for fibrillogenesis, lead to the formation of recombinant protein fibrils morphological similar to the parental seed. The results of the work can be used both in studying the fundamental mechanisms of conformation transfer and in developing test systems for synucleinopathies.

## Introduction

Synuclein is a protein with a molecular weight of 14460 Da, participating in synaptic transmission. Abnormal functioning of synuclein leads to the development of synucleinopathies. The role of synuclein aggregation in the development of Parkinson’s disease and amyotrophic lateral sclerosis has been previously shown [1]. In the literature, it has been discussed whether fibrils are capable of transmitting abnormal protein conformation from cell to cell - and whether synucleinopathies can be transmissible? A relatively recent study showed that fibrils isolated from patients with Parkinson disease (PD) and Amyotrophic lateral sclerosis (ALS) can be used for differential diagnosis of these diseases using the protein misfolding cyclic amplification (PMCA) and real-time quaking-induced conversion (RT-QuIC) assays [2], [3], [4]. In [5]using cryoelectron microscopy, it was shown that different types of synuclein fibrils differ in structure and, accordingly, have different ability to form complexes with fluorescent dyes and resistance to enzymatic hydrolysis by proteinase K, but such different fibrillar form features were not able to be transferred to the recombinant protein in native conformation [6]. One of the hypotheses explaining the formation of two significantly different fibrillar structures from one protein is the presence of different post-translational modifications (PTM) of synuclein in patients with different PD and ALS. In our study, we obtain two varieties of fibrils spontaneously formed from the recombinant protein that differ in morphology (according to transmission electron microscopy) and ability to increase the Thioflavin T fluorescence. The two fibril types obtained in the same conditions from the recombinant protein were able to transfer their morphological features and peculiarities in Thioflavin T tests when it was used as seeds. It is hypothesized that the difference in fibrils is associated with differences in the regions that determine the fibrillogenaseis during aggregate formation.

## 1. Materials and methods

### 1.1 Reagents

All reagents used were were manufactured by Sigma-Aldrich (USA) and Merck (USA).

### 1.2 Obtaining recombinant synuclein

In the first step, we inserted the yellow fluorescent protein (YFP) gene into the pET22b(+) plasmid (Novagen) as follows. Using restriction endonucleases BamHI / XhoI, a fragment (746 bp) containing the YFP gene was excised from the pTDpelB-C_sfYFPTwinStrep [7] plasmid. The excised fragment was inserted into the pET22b(+) plasmid treated with restriction endonucleases BamHI and SalI. The resulting plasmid pET22b(+)PelB-YFP-His was designed so that the resulting target protein would have the amino acid sequence (see Supplement, Figure 1). In the second stage, the obtained plasmid pET22b(+)PelB-YFP-His was treated with restriction endonucleases NdeI and BamHI, at the sites of which the alpha-synuclein gene was inserted. In this case, between the alpha-synuclein gene and the YFP gene, we placed a nucleotide sequence encoding a low-efficiency site for TEV protease [8]. Based on the obtained plasmid pET22b(+)alphasyn-TEV-YFP-His, a chimeric yellow fluorescent alpha-synuclein (alphaSyn-YFP) was synthesized, schematically depicted below:

**<alpha-synuclein>-<TEV protease recognition site>-<YFP>-<6xHis>-<stop>**

**Figure 1.**
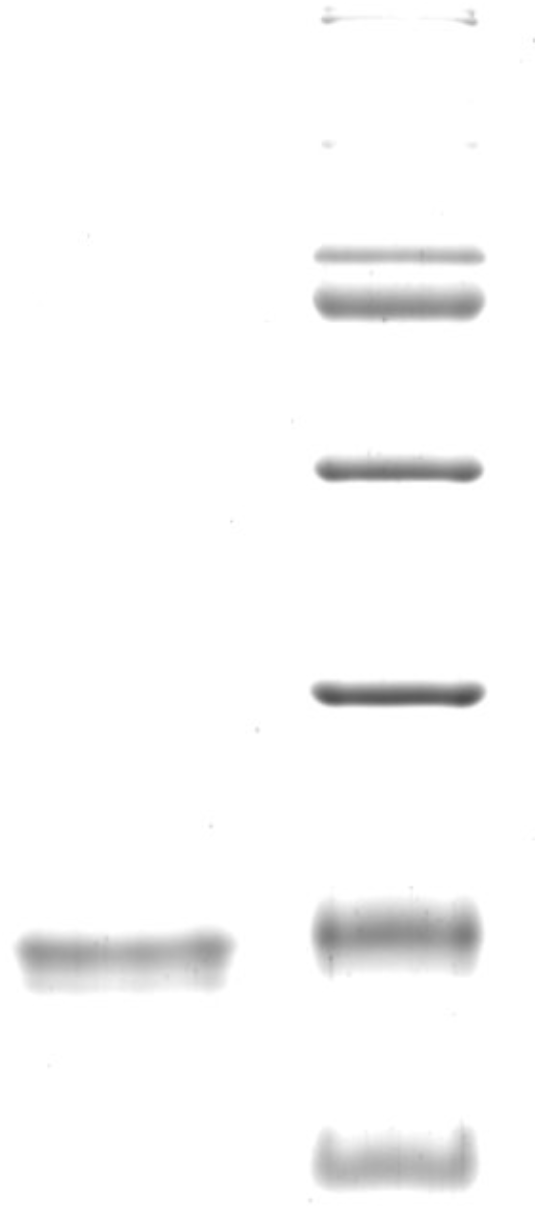
(A) Results of recombinant alpha-synuclein electrophoresis (B) results alpha-synuclein identification by mass-spectrometry. To reveal the ability of purified protein to form amyloid-like fibrils four identical probes of alpha-synuclein were incubated under fibrillogenic conditions (Materials and Methods 1.5). These probes were analyzed for their ability to bind to thioflavin T (Figure 2 C). It was found that there are two sorts of fibrils formed, differing in their ability to increase fluorescence intensity of the thioflavin T. Significant differences between the fluorescence intensity of thioflavin T were detected (Figure 2 A). A sample with an average level of fluorescence was also observed (F1), but in further studies we analyzed the properties the extreme cases (samples F2 and F4). Electron microscopy of the obtained fibrils F2 (Figure 2 A), showing the highest level of thioflavin T fluorescence, and F4 (Figure 2 B), showing the low level of thioflavin T fluorescence, was carried out. It turned out that the morphology of the fibrils were different. F2 sample shows presence of thin unbranched fibrils, at the same time in F4 sample fibrils, on the contrary, were stuck together and intertwined.

All the above components were in the same reading frame with the ATG start codon following the ribosome binding site (see annotated sequence at Supplement, Figure 2).

**Figure 2.**
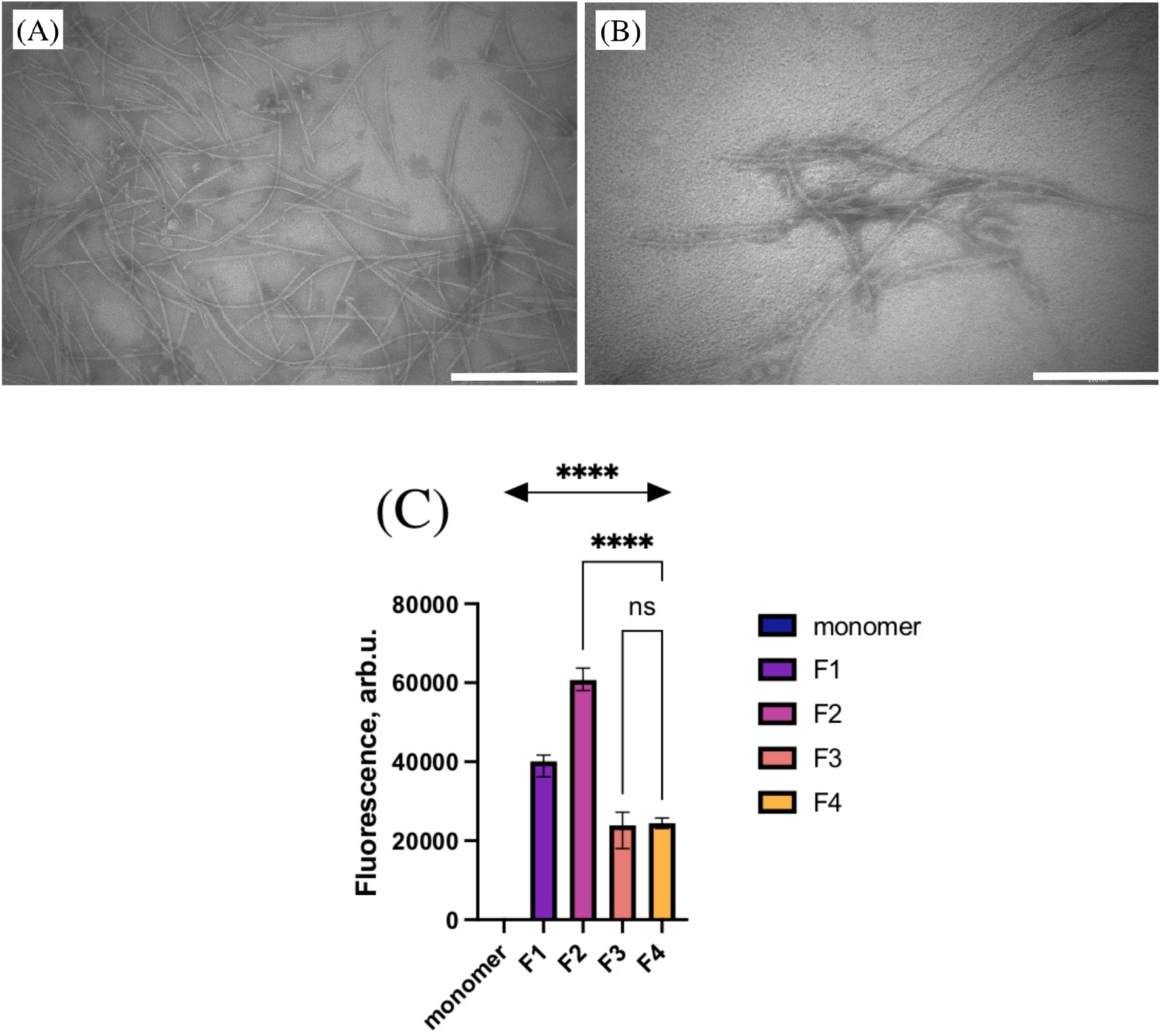
Spontaneous formation of different types of alpha-synuclein fibrils detected by TEM and ThT fluorometry: (A) and (B) electron microscopy of F2 and F4 samples; (C) fluorimetry intensity of ThT in presence of different samples. The length of the rectangle in the lower right corner corresponds to 200 nm.

Competent *E. coli* BL(DE3) cells were transformed with pET22b(+)alphasyn-TEV-YFP-His. The target protein synthesis was induced by a low concentration of IPTG (84 μM) for 20 h under very vigorous shaking (37°C). Yellow fluorescent alphasyn-YFP protein was detected both in the culture medium and in the periplasm. It was easily separated on a Ni-agarose column using standard techniques. The buffer for loading the affinity column consisted of 0.15 M NaCl, 10 mM imidazole.

In the third step, we performed dialysis against PBS with subsequent treatment of alphasyn-YFP with TEV protease, which we obtained in the method described in [8]. The efficiency of specific proteolysis was checked using 5-12% denaturing electrophoresis. Upon achieving the desired proteolysis result, the reaction mixture was purified on the same Ni-agarose column. TEV protease had 7 histidine residues at the N-terminus, and the cleaved YFP had 6 histidine residues at the C-terminus. To more effectively remove impurities containing oligohistidine sequences, the loading buffer was this time “softer” and consisted of 0.3 M NaCl, 7 mM imidazole. Fractions that did not bind to the column contained purified alpha-synuclein. The yield of pure alpha-synuclein was, on average, 80 mg per liter of culture medium. After dialysis against 10 mM Tris-HCl, pH 7.5, the protein was frozen at -80°C for long-term storage.

### 1.3 Polyacrylamide Gel Electrophoresis (PAGE)

Denaturing electrophoresis was performed according to the method [9]5 μl of a solution containing 2% SDS, 2.5% beta-mercaptoethanol and 0.01% bromophenol blue were added to 5 μl of protein at a concentration of about 1 mg/ml, then samples were incubated in a boiling water bath for 3 minutes. Protein electrophoresis in 12% PAAG was performed in 10×10 cm plates at a voltage gradient of 20 V/cm. The gel was stained with Coomassie R-250 solution according to [10]; images of the stained gel were obtained in a ChemiDoc MP gel-documenting station (BioRad, USA).

### 1.4 Mass spectrometry

To analyze the amino acid sequence of the protein after PAGE, enzymatic hydrolysis in the gel with trypsin was performed. A fragment of the stained zone was cut out, washed from the dye (twice with 100 μl of a solution of 30 mM ammonium bicarbonate, 40% acetonitrile in water), dehydrated in 100% acetonitrile, after which acetonitrile was collected and the gel fragment was incubated in air until complete evaporation of acetonitrile. Then 2 μl of trypsin solution (20 μg/ml in 50 mM ammonium bicarbonate) were added to the gel fragment and incubated in a solid-state thermostat GNOM (“DNK-Tecknologii”, Russia) at 37°C for 18 hours. The reaction was stopped with 3 μl of a solution of 1% TFA, 10% acetonitrile in water. To identify proteins, the resulting set of tryptic peptides was mixed with the DHB matrix in equal volumes, applied to a steel target and analyzed in the reflective mode of positive ion registration on an UltrafleXtreme (Bruker, Germany) MALDI-TOF/TOF mass spectrometer. At least 5000 laser pulses were summed for each spectrum.

Protein identification was performed using MASCOT [11]by simultaneously accessing the SwissProt database [12]and the local database, where the sequence of the synuclein recombinant protein was added. The accuracy of mass determination was limited to 50 ppm. Up to two trypsin errors (missing a proteolytic site) were allowed.

### 1.5 Fibril formation

Fibrillogenesis was carried out by incubation at 37°C with shaking at 500 rpm for 7 days in thermo shaker (Biosan TS-100); alpha-synuclein concentration was 0.4 mg/ml. The efficiency of fibrillogenesis was checked by analyzing the increase in fluorescence of the fibril complex with thioflavin T, as well as by analyzing the shift in the absorption spectrum of the fibril complexes and Congo Red. We observed the most pronounced fibrillogenesis when alpha-synuclein was incubated under the following conditions: 100 mM NaAc pH 3.0, 10 mM Tris-HCl pH 7.5, 25 mM NaCl, 0.05% NaN3 For seeding experiments, sonication of formed fibrils was performed in an ultrasonic bath for 15 minutes; 1/100 of the sample volume was added to the native protein solution in the same buffer.

### 1.6 Transmission electron microscopy

Electron microscopic analysis was performed to characterize the presence, shape and size of the synuclein protein fibrills. Samples were prepared for negative contrasting using 2% phosphotungstic acid solution. The sample (13 µl) was applied to a piece of parafilm (Parafilm M, Pecheney Plastics Packing, USA) fixed in a Petri dish, a 200-cell copper microscopy grid (Electron Microscopy Science (EMS), USA) with a carbon coating was placed on top of the sample, and incubated for 2 minutes. After that, the sample was removed and washed twice with 20 µl of distilled water for 15 seconds. The water was removed, the remaining moisture was blotted with a paper filter, and the applied preparation was contrasted with a 2% aqueous solution of sodium phosphotungstic acid for 2 minutes. After the time had elapsed, the grid was dried at room temperature for about 5 minutes. The measurements were carried out on a JEOL JEM 1100 electron microscope (JEOL, Japan) with an accelerating voltage of 80 kV.

### 1.7 Fluorimetry

20 μl of the sample were added to 80 μl of the buffer containing thioflavin T (to a concentration of 10 μM thioflavin T), thoroughly mixed and incubated for 15 minutes in the dark. Measurements were performed in 96-well darkened NUNC plates using a BMG Clariostar plate spectrofluorometer in fluorescence mode with an excitation wavelength of 440-10 nm, fluorescence was recorded at 478-10 nm. The signal gain value was set based on the fluorescence maximum in the studied series of samples. The signal was recorded along the well bottom using integration over the entire area of the well. The results of at least three measurements were used for processing. The SPSS SigmaPlot program was used to process and visualize the results.

## 2 Results

Recombinant synuclein was obtained as described in Materials and methods section. The protein fraction was subjected to electrophoresis. The protein purity was more than 90 percent. The structure was confirmed by mass spectrometry. The electropherogram is shown in Figure 1, and mass-spectrum is shown in the Supplement, Figure 3.

**Figure 3.**
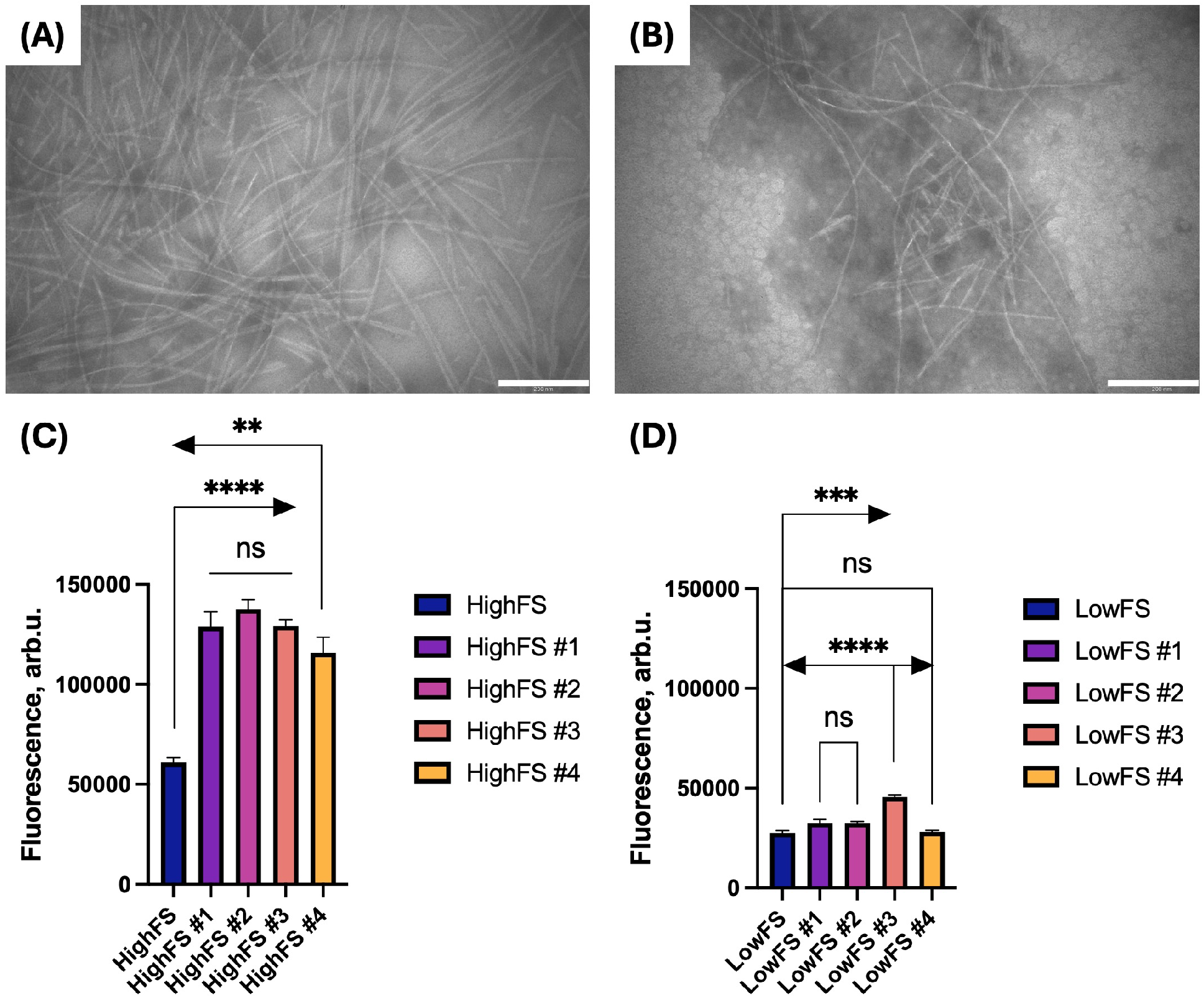
Induced formation of 2 types of alpha-synuclein fibrils detected by TEM and fluorometry. The length of the rectangle in the lower right corner corresponds to 200 nm.

The obtained signifantly different types of fibrils F2 (named «high fluorescence seed») and F4 (named «low fluorescence seed») were used as a seed for the growth of recombinant alpha-synuclein fibrils. The fibrils obtained using the seed also showed different ability to bind to Thioflavin T, and had the morphology similar to the parental fibrils (Figure 3 A and B, samples “HighFS#2” and “LowFS#2” were used for TEM).

It should be noted that the characteristic features of fibril morphology observed in seed samples are preserved during fibril growth from recombinant protein after seeding. Regarding the “HighFS” samples, it can be noted that the introduction of a high-fluorescence seed may allow synuclein to acquire a conformation that promotes more intensive fibril growth (all 4 samples have a higher fluorescence level with ThT compared to the seed, reliably). Moreover, the addition of the seed leads to more uniform growth (3 out of 4 samples do not differ from each other). In the “HighFS” samples, the addition of such a seed does not lead, unlike “HighFS”, to uniform growth of fibrils between biological replicates, which, although similar to the features of fibril growth without a seed, however, unlike growth without a seed, no samples are observed that have a high level of fluorescence (all of them are less than seeded by “HighFS”), that is, in this case, the seed affects growth, and apparently prevents the growth of fibrils with a high level of fluorescence. Thus, it was shown that the original recombinant protein can form at least two different types of fibrils. Morphology peculiarities and ability to form complexes with Thioflavin T can be transferred to new fibrils when using the obtained ones as a seed. The results allow us to suppose that the significantly different transferred conformation of alpha-synuclein may arise not as a result of PTM, but as a result of the ability of alpha-synuclein to spontaneously form different fibrils.

## Discussion

It has been shown that different types of fibrils are formed in ALS and Parkinson’s disease [2]. Moreover, when used as a seed in the protein misfolding cyclic amplification test, these different types of fibrils also initiate the formation of different types of fibrils. This can be explained both by the presence of post-translational modifications in proteins isolated from patients and by a significant difference in the synuclein conformations that arise in these pathologies. In [13], [14] it was demonstrated that the capacity of forming at least four types of fibrils from recombinant alpha-synuclein and described the conditions (variable buffer composition) under which one or another type of fibrils is formed. Moreover, the protein structures in these different types of fibrils were obtained using cryoelectron microscopy and an explanation is given for the observed effect of the buffer composition on the contacts between monomers [15], [16]. In this study, on the one hand, we use only recombinant proteins - and under the same conditions we obtain two types of fibrils that differ both in their ability to change the intensity of thioflavin T fluorescence and morphologically, according to transmission electron microscopy. Thus, we show that not only environmental factors and post-translational modification of proteins in fibrils used as a primer, but also simply the conformational lability of synuclein can lead to the formation of significantly different fibrils capable of transferring properties by a prion-like mechanism. In the future, the results of the work can be used both in studying the fundamental mechanisms of conformation transfer and in developing test systems based on synuclein fusion proteins with yellow fluorescent protein, for example, for diagnostics based on changes in fluorescence lifetime [17].

## Acknowledgments

The research was supported by RSF (project No. 23-24-00586)

## Supplement

**Figure 1.**
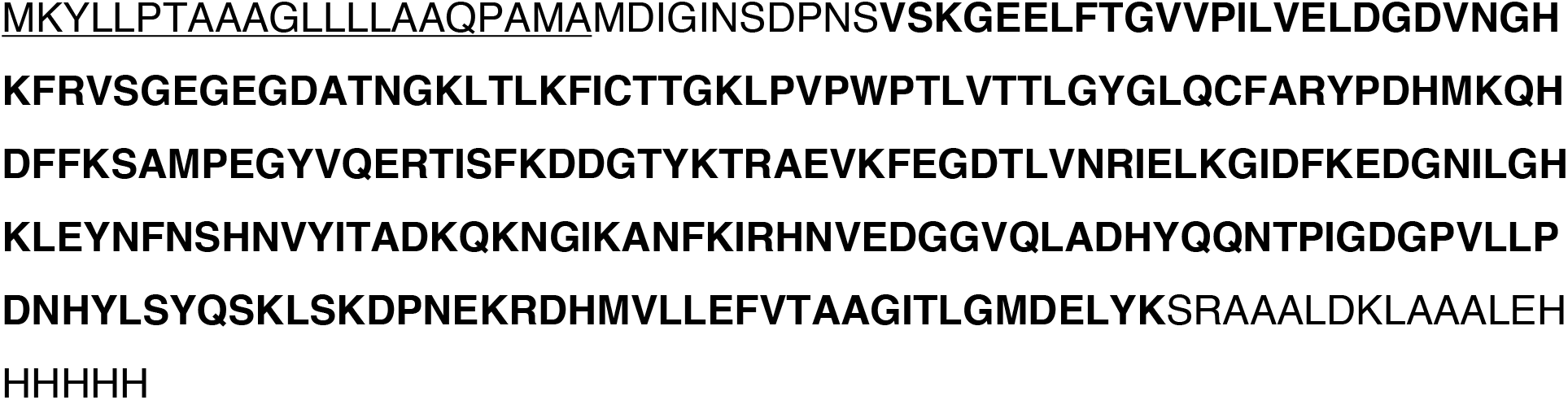
Recombinant YFP protein sequence. PelB leader peptide for export of target protein to *E*.*coli* periplasmic space is underlined. Oligohistidine sequence at C-terminus for isolation on Ni-agarose. YFP amino acids are shown in bold.

**Figure 2.**
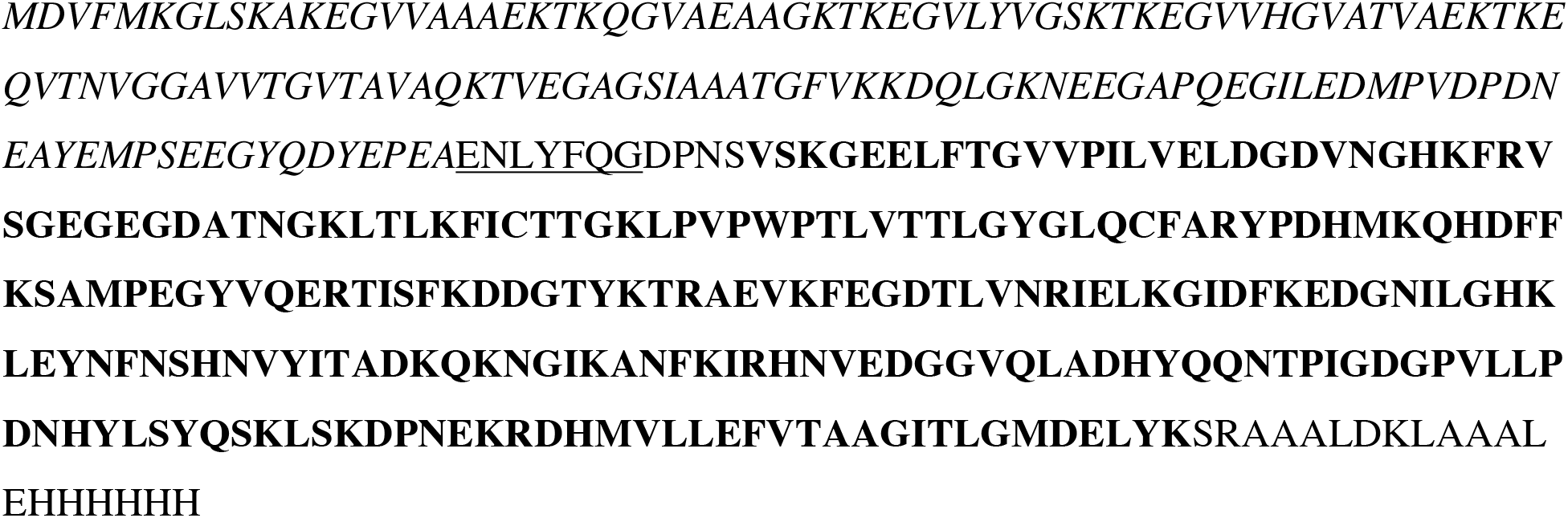
Recombinant YFP-synuclein fusion protein sequence. The amino acid sequence of alpha-synuclein is shown in italics. The TEV protease recognition site is underlined. The YFP amino acids are shown in bold. At the C-terminus is an oligohistidine sequence for highlighting on Ni-agarose.

**Figure 3.**
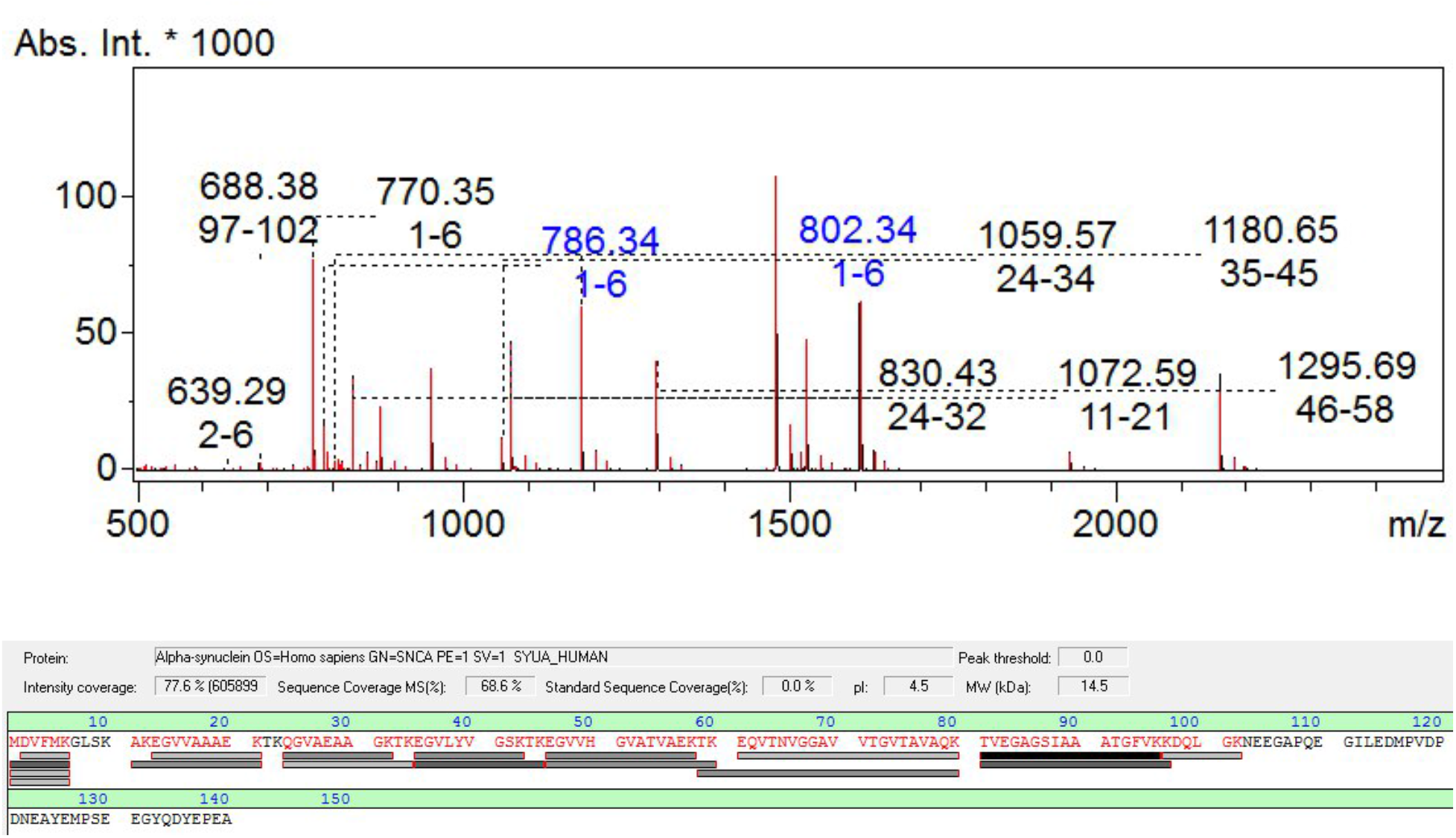
Results of alpha-synuclein identification by mass-spectrometry.

